# Landscape and dynamics of transcription initiation in the malaria parasite *Plasmodium falciparum*

**DOI:** 10.1101/024356

**Authors:** Sophie H. Adjalley, Christophe D. Chabbert, Bernd Klaus, Vicent Pelechano, Lars M. Steinmetz

## Abstract

The lack of a comprehensive map of transcription start sites (TSS) across the highly AT-rich genome of *P. falciparum* has hindered progress towards deciphering the molecular mechanisms that underly the timely regulation of gene expression in this malaria parasite. Using high-throughput sequencing technologies, we generated a comprehensive atlas of transcription initiation events at single nucleotide-resolution during the parasite intra-erythrocytic developmental cycle. This detailed analysis of TSS usage enabled us to define architectural features of plasmodial promoters. We demonstrate that TSS selection and strength are constrained by local nucleotide composition. Furthermore, we provide evidence for coordinate and stage-specific TSS usage from distinct sites within the same transcriptional unit, thereby producing transcript isoforms, a subset of which are developmentally regulated. This work offers a framework for further investigations into the interactions between genomic sequences and regulatory factors governing the complex transcriptional program of this major human pathogen.

## Introduction

Elucidation of the 23-Mb genome sequence of *P. falciparum*, the protozoan parasite responsible for the most lethal form of human malaria, revealed that its global AT-content is greater than 80% and rises to 90% in introns and intergenic regions (Gardner *et al*, 2002). This extreme base composition raised questions on the genome organization and mechanisms controlling gene expression in such an AT-rich environment. Transcriptome analyses using microarrays and later confirmed by RNA-Seq methods showed that gene expression is tightly regulated during the parasite intra-erythrocytic developmental cycle (IDC), which is associated with most of the disease symptoms (Bozdech *et al*, 2003b; Le Roch, 2004; Otto *et al*, 2010; Sorber *et al*, 2011). Indeed, the majority of the ∼5500 genes vary significantly in steady-state mRNA levels between the different intra-erythrocytic stages, resulting in a developmentally-linked cascade of gene expression. Genome surveys showed conservation of the basal eukaryotic transcriptional machinery, but also suggested a paucity of transcription-associated factors (Bischoff & Vaquero, 2010; Callebaut *et al*, 2005; Coulson *et al*, 2004). Despite major advances comprising the identification of transcription factors that govern *P. falciparum* stage-specific gene expression (Balaji, 2005; Campbell *et al*, 2010; De Silva *et al*, 2008), the lack of a comprehensive map of transcription initiation events has hampered advances in decrypting the molecular basis of transcriptional control. For instance, the mechanisms enabling the transcription machinery to cope with the exceptionally AT-biased nucleotide composition of the parasite genome or the basic organization of transcriptional units have yet to be determined. More particularly, the processes underlying the regulation of transcription initiation, such as the recruitment of RNA polymerase II (RNA PolII) or transcription start site (TSS) selection, still remain to be deciphered. Generally, characterizing the transcriptome architecture of *P. falciparum* has been technically challenging, given the low-complexity of its genome and the potential for reverse-transcription-derived artifacts (Siegel *et al*, 2014). Nevertheless, several single gene studies have attempted to identify transcription initiation sites (Horrocks *et al*, 2009), while the FULL-Malaria project mapped TSS positions for ∼25% of *P. falciparum* genes by cloning and subsequently sequencing full-length cDNA molecules (Watanabe *et al*, 2001; 2002). However, the low coverage of this single time point study did not address the intricacy and dynamics of transcription initiation in this parasite. The establishment of next-generation sequencing technologies now provides powerful means for the comprehensive and systematic analysis of gene expression and regulation. Genome-wide approaches such as CAGE (cap analysis of gene expression) have contributed to a better understanding of eukaryotic promoter structures and the impact of local nucleotide content on transcriptional processes (Shiraki *et al*, 2003). Recent characterization of the eukaryotic landscape of transcription initiation by RNA PolII has, for instance, revealed the existence of “broad” and “sharp” promoter classes (Carninci *et al*, 2006; Lenhard *et al*, 2012; Zhang, 2005) and shed some light on the mechanisms underlying the selection of transcription start sites (TSS) (Haberle *et al*, 2014).

Here, we report an in-depth analysis of the dynamics of transcription initiation during the *P. falciparum* IDC, giving new insights into the transcriptome architecture and gene expression regulation in this parasite. Using 5’ cap sequencing, a modified version of the CAGE approach that alleviates some of the artifacts previously mentioned, we systematically characterized *P. falciparum* transcript 5’ ends at different stages of the 48-hour intra-erythrocytic cycle, thereby generating a genome-wide TSS map at single nucleotide-resolution. We demonstrated that transcription initiation occurs across broad genomic regions and that multiple promoters may co-exist within a gene locus to generate mRNA isoforms. Strikingly, examination of the temporal data revealed that transcription initiation from separate sites within the same transcriptional unit may be coordinated or stage-specific, leading to the production of developmentally regulated transcript isoforms for a subset of *P. falciparum* genes. Analysis of the nucleotide content and chromatin organization around the newly defined TSS allowed us to define features of *P. falciparum* promoter architecture.

In particular, we found that even in the parasite’s extremely AT-biased genome, promoter selection and activity are governed by local base composition. Given the great complexity of the *P. falciparum* transcriptional program, this reports constitutes a highly valuable resource for further investigations into the mechanisms directing TSS selection, including the links between *P. falciparum* promoter architecture and regulatory elements. Our study also provides a framework for the development of novel therapeutics interfering with gene expression regulation in this deadly pathogen, to which half of the world population remains exposed (World Health Organization, 2014).

## Results

### Characterization of *P. falciparum* transcription initiation patterns reveals TSS distributions over broad promoter regions

To conduct the in-depth survey of the transcription initiation events occurring across *P. falciparum* developmental stages, RNA was extracted from two tightly-synchronized biological replicate cultures. These were harvested at 6 time points during the 48-hour IDC to examine ring (2 hours post-invasion (hpi), 10-hpi), trophozoite (18- and 26-hpi), and schizont stages (34- and 42-hpi).

Strand-specific libraries for deep-sequencing were constructed from the 5’ ends of capped mRNAs captured using a biotinylated 5’ adapter containing unique molecular identifiers, as reported in (Pelechano *et al*, 2015). In addition to CIP and TAP treatments, enrichment for capped mRNA molecules was further ensured by enzymatic treatment of the total RNA samples with a 5’ P-dependent exonuclease (Supplementary Information). Some modifications were made to take into consideration the exceptional AT-richness of the *P. falciparum* genome (Fig 1A, Supplementary Figs S1A-B, Material and Methods and Supplementary Information). For instance, the mRNA fragmentation and the reverse transcription reaction were performed using conditions previously established for RNA-seq and DNA microarrays in *P. falciparum* (Hoeijmakers *et al*, 2013a; Bozdech *et al*, 2003a). Additionally, the second strand synthesis as well as the amplification of the cDNA libraries were performed using the KAPA DNA polymerase enzyme that has a very low AT-bias and is now commonly used when preparing *P. falciparum* sequencing libraries (Oyola *et al*, 2012; Siegel *et al*, 2014). About 600,000 unique 5’ long tags (> 90 nucleotides) were recovered on average per independent library (Supplementary Fig S1C, Supplementary Table S1) and collapsed on their first base to define the position of the transcription start sites (TSSs) at single-nucleotide resolution. .Compilation of the transcription initiation events detected on both positive and negative strands showed that more than 3 million nucleotide bases constituted a TSS, indicative of an extensive transcription initiation activity across the *P. falciparum* genome (Supplementary Table S1). The use of in vitro transcripts to control for the robustness of the protocol confirmed that the 5’ end of these transcripts were properly identified using the 5’ cap capture (Supplementary Fig S1D, Supplementary Information).

**Figure 1.**
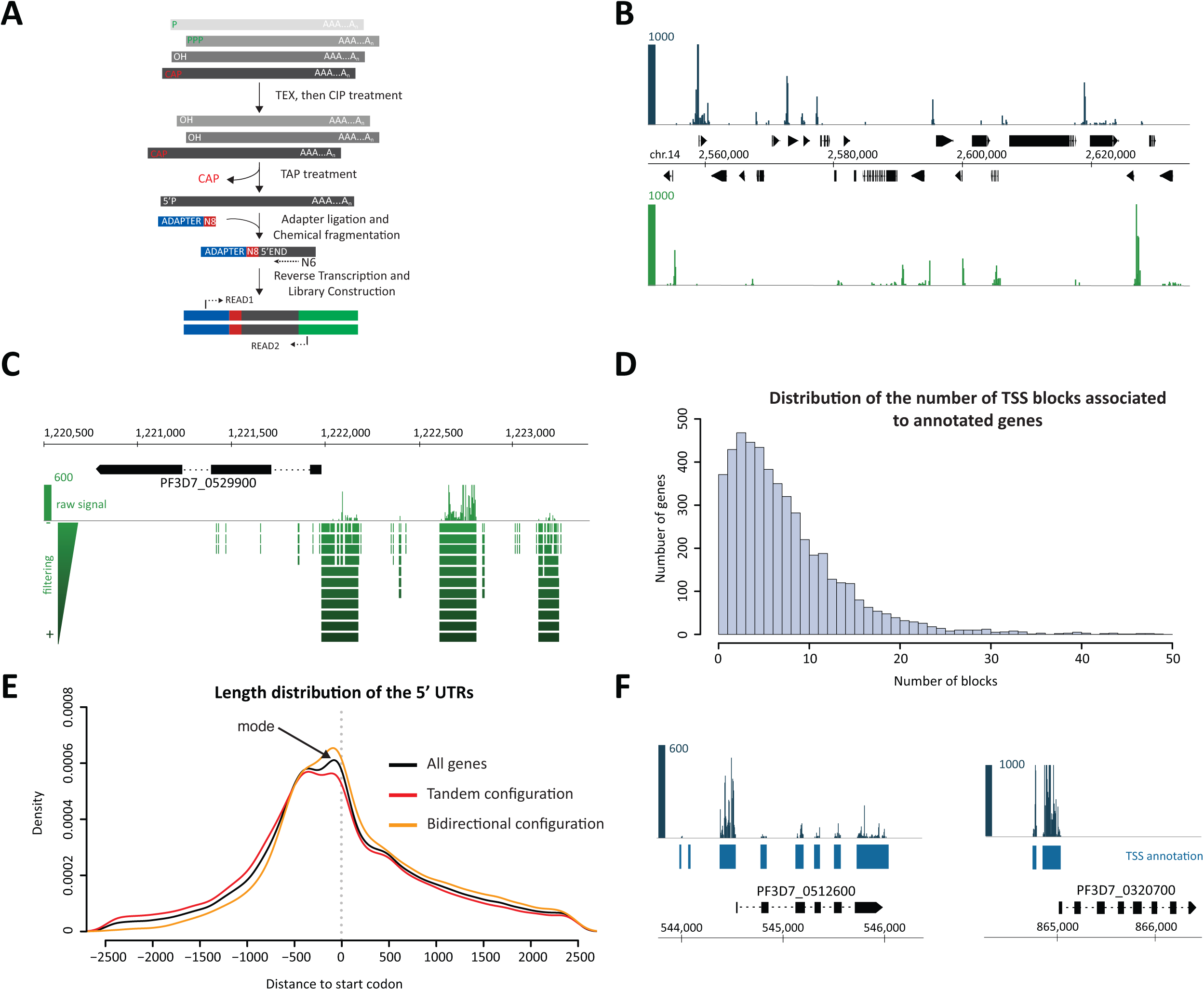
Genome-wide identification of TSS during *P. falciparum* asexual cycle. (A)Capped mRNA molecules are captured using direct ligation of RNA adapters containing a molecular barcode to transcript 5’ ends. Enrichment for capped mRNA molecules is achieved by treating total RNA samples with a 5’ P-dependent exonuclease (TEX), followed by a treatment with an alkaline phosphatase (CIP). Illumina libraries are then amplified after reverse transcription of the tagged mRNA fragments. (B) Snapshot of a coverage track showing the intensity of the TSS mapped to a locus on chromosome 14. Filtered reads from all time points and biological replicates were pooled and collapsed on their first base pair to visualize sites of initiation only. The scale indicates the read counts at each genomic position. (C) Snapshot of a coverage track illustrating the annotation obtained using alternate sequencing filters. Read counts show the profile of the raw signal. Green boxes show the annotation of TSS blocks generated at each step of the filter size increments. (D) Distribution of the number of TSS blocks associated with previously annotated transcriptional units. (E) Length distribution of the 5’ UTRs as obtained from our new annotation. While most TSS are located upstream of the start codon, a large number of transcripts have very short (<100bp) or no 5’ UTRs due to initiation within gene bodies (black). Length distribution of the 5’ UTRs for genes in a tandem (red) and bi-directional configuration (orange) is also shown. (F) Coverage tracks showing examples of the distribution of TSS across the body of multi-exons genes. Transcription widely initiates within the body of each exon of PF3D7_0512600, whereas TSS clusters are only detected upstream of the start codon for PF3D7_0320700. TSS block annotation is indicated in blue.

Transcription initiation events in *P. falciparum* appeared to occur at multiple closely spaced sites, across relatively wide regions (Fig 1B), indicative of a broad promoter architecture similarly to other eukaryotic organisms (Lenhard *et al*, 2012). We therefore implemented an analytical method that adapts to the various types of captured signals (dispersed or dense distribution, for instance) and allows to define separate promoter regions or “TSS blocks” in a systematic way, instead of clustering the TSSs within an empirically chosen window size across the genome (Fig 1C). Mathematical morphology operations and filtering tools (Heijmans *et al*, 1989) were applied to the pooled data to identify all separate blocks of transcription initiation independently of the replicates and developmental stages (Material and Methods). The use of alternating sequential filters led to the definition of more than 44,000 potential TSS blocks, thus promoter regions, genome-wide that were associated where possible to one of the annotated transcriptional units (Supplementary Fig S2 and Material and Methods) to constitute a high-resolution atlas of TSSs for the *P. falciparum* transcriptome. Biological replicates exhibited an excellent correlation (Spearman correlation > 0.9), thereby confirming the robustness of our protocol (Supplementary Fig S1E). In total, between the two replicates, transcription initiation events were detected for 90% of the *P. falciparum* annotated genes (4,955 out of 5,510). The vast majority (92%) of these contained more than one cluster of transcription initiation sites (median of ∼ 6 TSS blocks), thereby expanding on previous observations from the partial analysis of full-length cDNAs by Watanabe et al in 2002 (Watanabe *et al*, 2002). This striking finding implies that the majority of the genes use not only multiple sites as evidenced by the dispersed patterns of transcription initiation, but also multiple promoter regions to initiate transcription (Fig 1D). Importantly, this confirms the additional level of resolution provided by our approach in comparison to that of current RNA-Seq technologies.

### Transcription in *P. falciparum* may initiate close to the coding sequence of genes

Given that the majority of genes appeared to use multiple sites to initiate transcription, we examined their global distribution relative to the coding region of the associated genes. This analysis showed that 81% of the total TSS blocks were positioned less than 1000-bp upstream of the start codon, and more than 65% of these located within a distance of 500-bp or less upstream of the CDS (Fig 1E). Interestingly, the mode of the TSS block distribution around the start of coding sequences (CDS) was within 50-bp upstream of the start codon (Fig 1E). This suggests that transcription tends to initiate in regions very close to the start of the CDS to produce leader-less transcripts, i.e transcripts with little or no 5’ UTR.

Our analysis further identified transcription initiation in non-conventional sites, such as within the coding region of single exon genes, but also in exons of multi-exon genes (Fig 1F). Individual instances of such configurations were also observed in the FULL-malaria dataset (Supplementary Fig S3). 49% of all identified TSS blocks were actually located downstream of the start codon. This indicates a significant amount of transcription initiation events within the CDS that potentially leads to truncated protein products as observed in yeast (Fournier *et al*, 2012). The number of internal TSS blocks that are followed by an ATG codon in frame with the annotated coding sequence (∼ 90%) suggests that it might also be the case in *P. falciparum*. However, some of the sequencing tags recovered in these regions may also result from recapping of mRNA degradation products or non-coding RNAs (Schoenberg & Maquat, 2009).

Further analysis showed that the distribution of the TSS blocks was similar whether genes displayed a tandem or bidirectional configuration (Fig 1E) and was not markedly affected by the presence of introns (Supplementary Fig S4A). Increased length of the intergenic region was associated with a wider spread of the TSSs upstream of the start codon, with the TSSs closer to the CDS for genes separated by a short distance (Supplementary Fig S4B). Annotated features for which no TSS was identified corresponded for the vast majority to antigenic variant families and to genes that are expressed in the other stages of the parasite infectious cycle (Supplementary Table S2A).

### *P. falciparum* transcriptome exhibits widespread intergenic and bidirectional promoter activities

An appreciable number of TSS blocks (4,600, i.e. 10%, across all 14 chromosomes) could not be assigned to any of the gene loci found in the current annotation of the *P. falciparum* genome. These were therefore categorized as “new blocks of transcription initiation” and manually clustered (Supplementary Fig S4C, Material and Methods) to isolate more than 1,500 “potential transcriptional units” (Fig 2A, Supplementary Table S2B). Most of these clusters are located in non-coding regions, or within very large intergenic regions between annotated genes, suggesting that a large fraction of the *P. falciparum* non-coding genome is actively transcribed (Mourier *et al*, 2008; Raabe *et al*, 2010; Wei *et al*, 2014). Such non-coding transcription could play a role in regulating gene expression, for instance via recruitment of chromatin modifiers (Fatica & Bozzoni, 2013) or by strengthening transcriptional activities through interactions with the transcriptional machinery (Barrandon *et al*, 2012). This postulate extends to the telomeric and subtelomeric regions of the parasite genome, in which a few non-coding transcripts had been detected and linked to transcriptional control and silencing (Broadbent *et al*, 2011; Epp *et al*, 2008; Kyes *et al*, 2007). We generalized these findings with the identification of new transcripts emerging from the telomeric regions of 11 out of all 14 chromosomes and within introns of at least 80% of all *var* genes (Supplementary Figs S4D and S4E, Supplementary Tables S3A and S3B).

**Figure 2.**
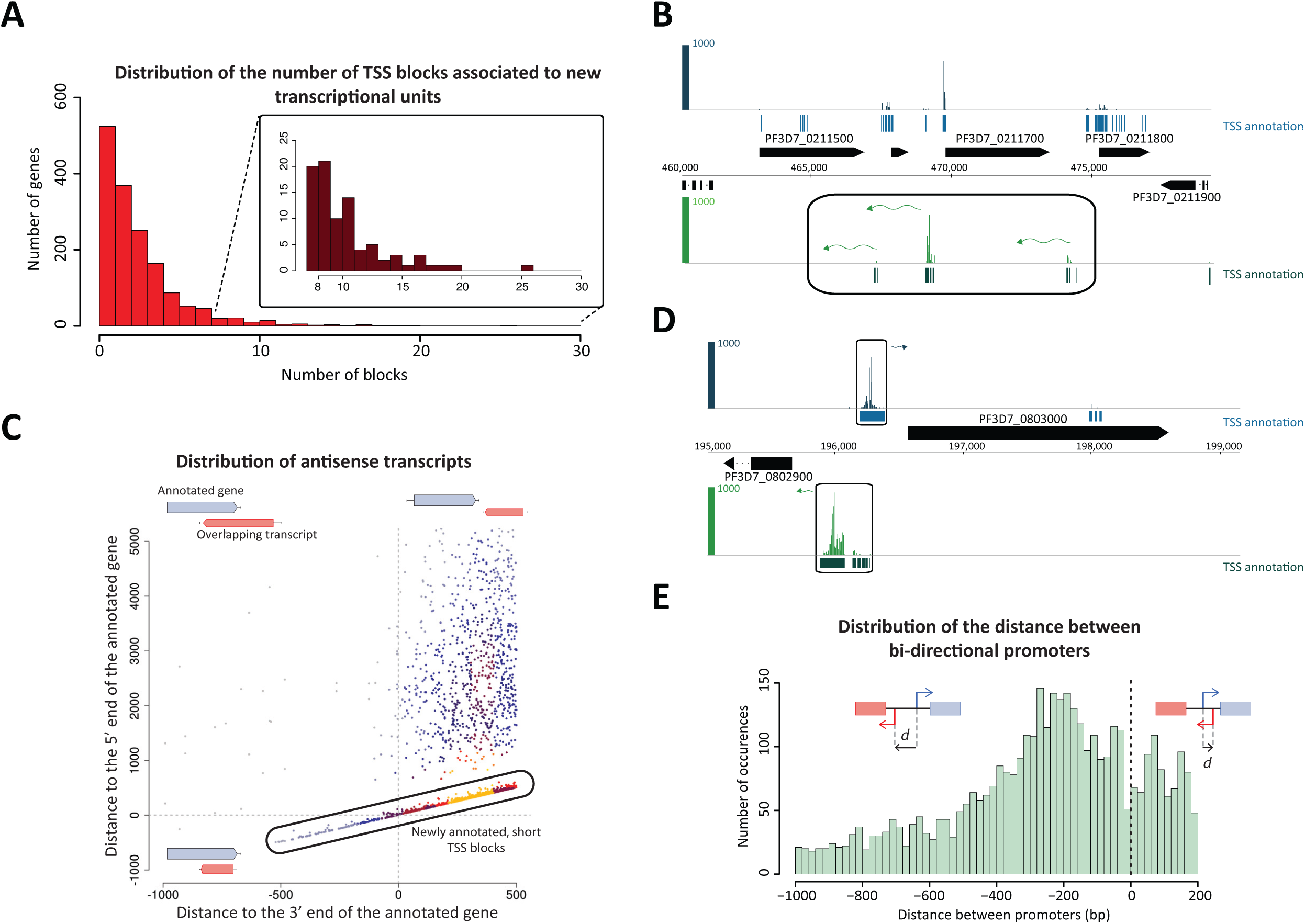
Global organization of the TSS along the *P. falciparum* genome. (A) Distribution of the number of TSS blocks associated with the newly clustered transcriptional units. The enlargement illustrates the presence of transcriptional units with a high number of TSS blocks (>7 per unit). (B) Snapshot of a coverage track for a genomic locus containing several newly annotated antisense transcripts that potentially overlap with the 3’ end of coding genes located on the opposite strand. TSS block annotations are indicated in blue (positive strand) and green (negative strand). (C) Distribution of the various categories of antisense transcripts defined by the distance from the end of the antisense transcripts to the annotated 3’ end of the sense transcript against the distance from the start of the antisense transcript to the annotated 5’ end of the sense transcript. This distribution reveals that most of the detected antisense transcripts initiate close to the 3’ end of annotated genes. (D) Snapshot of a coverage track depicting a bidirectional promoter in which transcription initiates in opposite direction from sites located a few hundred base pairs apart. (E) Distribution of the distance between the 5’ end of bidirectional promoter elements. Negative distances correspond to TSS located upstream of the 5’ reference point while positive values correspond to TSS located downstream (and therefore potentially resulting in overlapping transcripts).

The pervasive aspect of *P. falciparum* transcription extended to antisense transcription initiation, which was detected for 31% of the 4,955 examined genes (Fig 2B). A large fraction (47%) of the ∼2,000 antisense TSS blocks identified genome-wide resulted from non-coding transcription. Interestingly, a third of these TSS blocks directly overlapped with the 3’ end of the sense annotated genes (Fig 2C), thereby plausibly interfering with the regulation of the derived sense transcripts. Further analysis showed that 33% of the antisense TSS blocks were located in promoter regions where transcription may initiate bidirectionally (Fig 2B). Such regions were prevalent, the presence of TSS blocks in a divergent arrangement being observed for about half of all annotated features (Fig 2D). In more than 70% of such regions, the pairs of divergent TSS blocks were separated by a distance of 400-bp or less from each other, arguing for the likely presence of bidirectional promoters with shared regulatory elements (Fig 2E). This type of configuration may allow for the co-expression of adjacent genes, or alternatively the directional control of gene expression through binding of regulators to separate promoter elements.

### Core promoter regions are defined by a specific chromatin organization

The generation of a genome-wide map of transcription initiation events now offers the possibility of assessing the structural properties that may define TSS choice in the malaria parasite. Among these, local chromatin organization plays an important role in the regulation of eukaryotic transcription initiation, with the association of specific histone marks and variants with active promoters. We therefore examined the dynamics of H3K4me3-, H3K9ac-, H2A.Z- and H2B.Z occupancies around the most active transcription start sites (see Material and Methods), using publicly available datasets of chromatin immuno-precipitation followed by sequencing (ChIP-Seq) (Bártfai *et al*, 2010; Hoeijmakers *et al*, 2013b) (Fig 3A).

**Figure 3.**
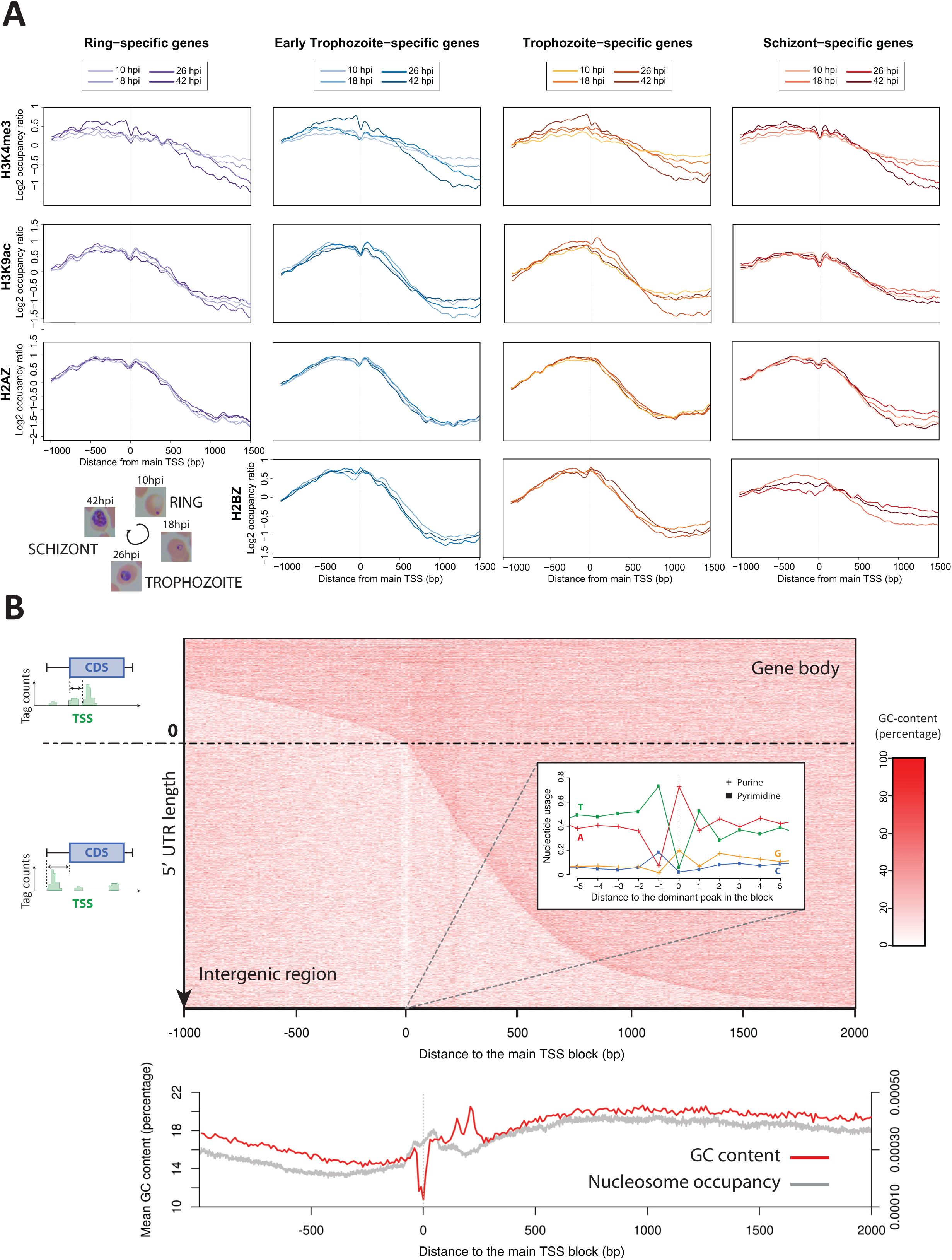
Architecture of active promoters during *P. falciparum* asexual cycle. (A) Occupancy of histone post-translational modifications and variants around stage-specific promoters. The occupancy profiles are organized by mark/variant (rows in the table) and stage-specific genes (columns in the table). In each case, the occupancy profiles were plotted for each time point at which the mark or variant was profiled (Bartfai et al. 2010, Hoeijmakers et al, 2013). Only the most active TSS block was considered for each gene and all ChIP signals were normalized using the chromatin input signal. Stage-specific promoters were called using the counts obtained from our study (See Supplementary Material). This global view illustrates developmentally regulated changes in the distribution of histone marks and variants around *P. falciparum* promoters. (B) Local nucleotide content and nucleosome occupancy around the most active TSS block. The heatmap illustrates the local GC content in 10-bp bins around the TSS block for each of the genes with an annotated TSS block. Analysis of the nucleotide composition within a symmetrical 10-bp window around the dominant TSS peak within the block shows the preferential pyrimidine (T)-purine (A) di-nucleotide used to start transcription. The average profile below the heatmap reveals the drop in GC content around the site of initiation as well as the +1 nucleosome barrier. The additional downstream peaks enriched in GC nucleotides are also apparent and delimit a decrease in nucleosome occupancy. Nucleosome occupancy was plotted using the Mnase-Seq data from (Bunnik *et al*, 2014).

When aligning all transcripts at the start position of the most active associated TSS block, we found predominant enrichment for these histone post-translational modifications and histone variants at the 5’ end of genes (Fig 3A), as in other eukaryotic species. H3K4me3- and H3K9ac-marked nucleosomes were precisely positioned at the +1 position, i.e. the first nucleosome downstream of the TSS block, suggesting that the strong relationship between TSS and position of the +1 nucleosome is maintained in the malaria parasite (Struhl & Segal, 2013). Nevertheless, both marks displayed highly discernible occupancy profiles. Globally, H3K4me3 enrichment increased steadily as the parasite progressed through the IDC, with a stronger marking of the +1 and +2 nucleosomes and a more pronounced nucleosome-free region (NFR) by the end of the cycle (Fig 3A). In contrast, H3K9ac enrichment at the +1 nucleosome position increased progressively before dropping at the schizont stage to a level equivalent or lower than that observed at early developmental stages (Fig 3A). Enrichment for the acetylation mark around the TSSs was more pronounced with a prominent NFR at the trophozoite stage, during which the bulk of transcriptional activity occurs (Sims et al., 2009).

Interestingly, the overall profiles of H3K4me3 enrichment were highly similar between groups of genes expressed at different developmental stages, confirming the disconnect between H3K4me3 occupancy and genes’ transcriptional activity (Fig 3A) (Bártfai *et al*, 2010). Genes specifically expressed late in the parasite IDC, on the other hand, exhibited very little variation in occupancy for H3K9ac-marked nucleosomes around the TSSs between time points (Fig 3A). This trend was also observed for nucleosomes containing the histone variants H2A.Z and H2B.Z (Fig 3A). Their occupancy profiles appeared constant and similar between genes expressed at distinct stages of the parasite life cycle, with a characteristic enrichment at the +1 nucleosome (Mavrich *et al*, 2008) and a stronger depletion towards the 3’xhibited expression profiles in opposite phase end of transcripts. In contrast, genes preferentially active during schizogony exhibited an increased enrichment for H2A.Z around the TSSs at the beginning and end of the life cycle, while distribution of H2B.Z was generally more even on either side of the TSS.

Altogether these observations indicate that although chromatin organization around the TSS displays expected features such as a marked positioning of the +1 nucleosome downstream of the TSS, the presence of the examined histone marks and variants do not tightly associate with transcriptional activation as observed in other eukaryotes.

### *P. falciparum* TSS are characterized by a particular nucleotide composition

In addition to examining the chromatin landscape around the sites of transcription initiation, the generation of such a high-resolution TSS map enabled us to assess whether *P. falciparum* core promoters are characterized by a particular nucleotide signature. Analysis of the base composition in the core promoter region after aligning all genomic sequences at the most active TSS block revealed a strong decline in GC content around the position of transcription initiation (Fig 3B). When looking at the dominant TSS peak within the block, we observed that transcription in the parasite preferentially initiates with the pyrimidine-purine di-nucleotide T-A at position -1, 0 (Fig 3B). This trend was independent of the activity of the considered TSS block (Supplementary Figs S5A and S5B). In contrast, no specific di-nucleotide composition was observed when randomly selecting a genomic position within the blocks (Supplementary Fig S5C). Surprisingly, we also detected the presence of two well-defined peaks corresponding to a local increase in G/C about 150-bp and 210-bp downstream of the TSSs, respectively. Alignment of the profile of GC-content and that of nucleosome positioning derived from (Bunnik *et al*, 2014) revealed that these two peaks coincided with a dip in nucleosome occupancy. Nucleotide composition around the TSSs did not vary strikingly with the strength of the promoter, except for a greater GC content right at the borders of the TSS blocks, and in particular downstream of those with a lower activity (i.e. less frequent usage) (Supplementary Figs S5D and S5E). This change was accompanied by a shift of the +1 nucleosome and a disappearance of the NFR around the TSSs, indicative of a direct link between nucleotide content, nucleosome positioning and transcriptional activity.

### Dynamic analysis of TSS usage during *P. falciparum* IDC reveals alternative transcription initiation and possibly shared regulatory elements

The generation of our genome-wide TSS annotation enabled us to examine the dynamics of transcription initiation during the *P. falciparum* IDC and identify significant differences in TSS usage between developmental stages. Stringent filters using the replicate information were applied (Supplementary Figs S6A-D) and confirmed extensive transcriptional activity during the *P. falciparum* IDC. In total, transcription initiation events conserved across replicates were detected for 74% of all 5,510 protein-coding genes (local FDR < 0.1), and 68% of all annotated features (4,838 out of 7,090, FDR < 0.1) that include the new transcriptional units as defined above.

Given that the *P. falciparum* IDC is characterized by a cascade of gene expression (Bozdech *et al*, 2003b; Llinas, 2006), we assessed whether this is recapitulated at the level of transcription initiation events. We found that the majority of the actively transcribed genome (91% of the 4,738 annotated features to which active TSS blocks were associated) exhibited a cycling behavior throughout the parasite developmental cycle (log2fold change > 0.5, Fig 4A, Supplementary Fig S6E). These numbers are in agreement with previous microarrays and RNA-Seq studies and confirm that the parasite transcriptional cascade is also observed at the TSS level.

**Figure 4.**
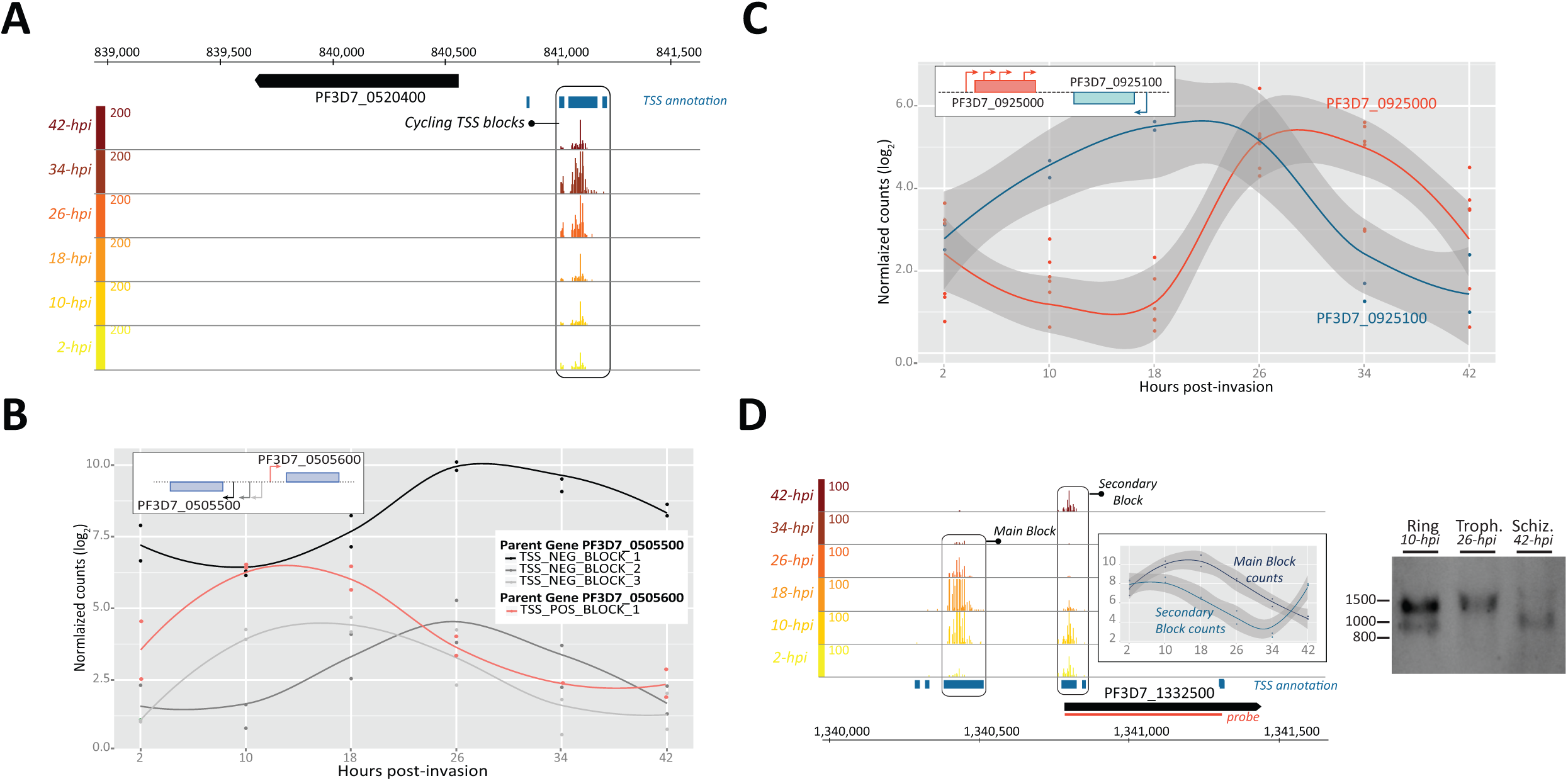
Dynamic usage of transcription initiation sites during *P. falciparum* intra-erythrocytic development. (A) Coverage tracks illustrating the periodic transcription initiation events associated with the gene PF3D7_0520400. This cycling transcriptional activity is characteristic of the reported cascade of gene expression during the parasite IDC. (B) Normalized counts of a set of asynchronous TSS blocks associated with a pair of bidirectional genes (PF3D7_0505500/PF3D7_0505600). The timing of maximal expression for the 4 blocks defines two classes of TSS clusters with different expression patterns and yet sometimes associated with the same gene. (C) Normalized counts of a pair of genes (PF3D7_0925000/PF3D7_0925100) in antisense orientation and displaying temporal usage of TSS blocks (marked as dots) in opposite phase. (D) Example of a gene, PF3D7_1332500, harboring two types of TSS blocks with different temporal usage during the IDC. The coverage tracks and normalized counts indicate a strong increase in activity at 18-hpi for the main TSS block, while the secondary block is mostly used at the early and late stages. Northern blot analysis clearly illustrates the changes in relative abundance of the two populations of RNA molecules (one being leader-less) between stages. The genomic location covered by the northern blot probe is highlighted in orange.

In view of the non-random organization of eukaryotic genomes, indicative of the functional importance of gene distribution (Hurst *et al*, 2004), we first investigated possible interdependency in TSS usage between genes depending on their arrangement in the genome. For the vast majority (91%) of genes pairs in divergent configuration, bidirectional transcription initiated coordinately, suggesting potential co-expression (Supplementary Fig S6F). However, a number of those (183 out of 2084, Supplementary Table S4A) exhibited distinct cycling behavior, as reflected by the contrasting profiles of the associated TSS blocks (Supplementary Fig S6G). Intriguingly, we also identified divergent gene pairs with distinct expression patterns that comprised TSS blocks with highly similar profiles of temporal usage (Fig 4B, Supplementary Fig S6H), suggesting the presence of both common and separate promoter elements. Altogether these results demonstrate that bidirectional promoters may contain multiple regulatory elements that would enable both coordinated and dissociated TSS usage within divergent transcriptional units. Given that many of the regions of bidirectional transcription were also sites of antisense transcription, we analyzed the temporal usage pattern of TSS blocks for cycling sense/antisense pairs (11% of all sense/antisense pairs, stringent local FDR threshold of 0.01). We included in our survey sense/antisense pairs on opposite strands in a convergent configuration. We observed dissociated transcription for most. Indeed, only a minority of the antisense transcription initiation events (<1%) occurred in synchrony with their sense counterparts, whereas a few pairs of sense/antisense transcriptional units (52 out of the ∼1100 pairs surveyed) exhibited expression profiles in opposite phase (Fig 4C, Supplementary Table S4B). These observations confirm that sense and antisense transcription initiation are sometimes interdependent, whereby antisense transcription acts as a regulator of gene expression, by promoting or preventing expression of the sense transcript (Wei *et al*, 2011; Werner, 2013).

Given that most transcriptional units in *P. falciparum* contain multiple TSS clusters, we additionally examined the dynamics of transcription initiation within each of those. For most of the annotated features, transcription appeared to initiate coordinately at each of the associated TSS blocks, suggesting that even when transcription starts at multiple sites within a given gene locus, all sites are concomitantly used (Fig 4A). Nonetheless, a subset of *P. falciparum* annotated features (124, 3.4%) exhibited TSS blocks with distinct cycling behavior (Fig 4D, Supplementary Table S4C). Among these, 57 harbored TSS blocks with temporal usage patterns in opposite phase, indicative of a switch between TSS in a stage-specific manner and suggesting that initiation events at these sites are mutually exclusive (Supplementary Table S4D). Northern blot analysis of selected candidates confirmed that transcription initiation at alternative sites does lead to production of full-length transcript isoforms (Fig 4D, Supplementary Fig S6I) whose expression is developmentally regulated.

## Discussion

This report constitutes the most extensive survey to date of the transcription initiation events occurring across various stages of the *P. falciparum* asexual blood cycle and provides insights into promoter architecture for a complex and thus far poorly characterized genome. Our data therefore represent a powerful resource that will enable further investigation into the molecular mechanisms governing the selection of TSS and more broadly transcriptional regulation in such an extremely base-biased environment.

Previous studies aiming to identify regulatory elements in the *P. falciparum* genome generally employed sequence motif search tools using genomic sequences at an arbitrarily chosen distance upstream of the translation initiation codon (Gunasekera *et al*, 2007; Militello *et al*, 2004; Young *et al*, 2008). Our genome-wide TSS map enables this type of analyses to henceforth directly focus on the genomic regions adjacent to the initiation sites to accurately distinguish between core promoter elements and binding sites for specific factors. The in silico survey of DNA physicochemical properties predicted core promoter regions using the partial TSS mapping from Watanabe et al. (Watanabe *et al*, 2002) and estimated thymine-adenine to be the preferred sequence at the TSS (Brick *et al*, 2008). We demonstrated that TSS selection at the genome-wide level is organized around such a specific di-nucleotide composition, reflecting the general preference for a pyrimidine-purine initiation site observed in other eukaryotes (Carninci *et al*, 2006). Given the low-complexity sequence context associated with the *P. falciparum* genome, the question therefore arises of what would define which genomic site is to be used for transcription initiation in such an AT-rich environment. We showed that the frequency of usage of a TSS is actually guided by the local G/C content at precise positions downstream of the TSS. This indicates that at least part of the structure of core promoters is genomically-encoded and that there might be some spatial constraints for the positioning of the basal transcription machinery. Indeed, the local increase in GC content around weaker TSS suggests the establishment of tighter boundaries for transcription initiation possibly by way of a more defined nucleosome positioning or particular epigenetic mechanisms such as DNA methylation (Ponts *et al*, 2013). In fact, our observation that the diminution in TSS strength is accompanied with an increase in the distance between the TSS and the +1 nucleosome together with a reduced NFR argues for an association between TSS selection, nucleosome positioning and regional nucleotide content. Altogether these observations indicate that despite the parasite’s extraordinary AT-biased genome, promoter selection and activity are primarily defined by local base composition and chromatin structure. Intriguingly, timing of TSS usage does not correlate with enrichment in the histone marks H3K4me3 and H3K9ac or the variants H2A.Z and H2B.Z at the 5’ end of genes, which are typically associated with promoter activity (Li *et al*, 2007; Zlatanova & Thakar, 2008). Instead these seem to be associated with *P. falciparum* maturation, independently of the activation of gene expression, in agreement with previous studies reporting a rather moderate association between the dynamic changes in the histone marks/variants and transcript levels (Bártfai *et al*, 2010; Gupta *et al*, 2013; Hoeijmakers *et al*, 2013b). Interestingly, the greatest changes in chromatin organization around the TSS occur at the schizont stage and for schizont-specific genes. This could reflect the replication activity that takes place during this developmental stage or a mechanism of synchronization of gene expression to prepare all future daughter cells for the next cycle.

Further examination of the dynamics of TSS usage captured the cascade of gene expression characteristic of the malaria parasite, whereby all TSS associated with the same transcriptional unit were coordinately used at a specific time during the parasite’s life cycle. For a vast majority of such transcriptional units, we observed a high recurrence of synchronous transcription initiation events from multiple sites. This may lead to the simultaneous production of transcript isoforms with alternative 5’ UTRs, as indicated by our northern blot analysis, and thus increased transcriptome diversity (Pal *et al*, 2011). The concomitant expression of several isoforms with varying 5’ UTRs may have important regulatory consequences (Davuluri *et al*, 2008), notably by influencing transcript stability (Hogan *et al*, 2008) or translation efficiency (de Klerk & Hoen, 2015). Interestingly, recent ribosome and polysome profiling studies in *P. falciparum* have reported enrichment for ribosomes along the 5’ UTR of numerous transcripts (Bunnik *et al*, 2013; Caro *et al*, 2014), including those lacking AUGs (Caro *et al*, 2014). This suggests the existence of upstream open reading frames (uORFs) that are actively translated from non-cognate initiation codons (Ingolia, 2014) and may affect the efficiency of translation of the full-length transcript. In addition to the numerous transcriptional events detected upstream of the translation start codon, we noticed that most of the transcriptionally active loci contained TSS that overlap with the start of the coding sequence. Our observations suggest that these events produce at least in some cases leaderless transcripts that may be translated (Cortes *et al*, 2013). The high number of multi-exonic genes containing exonic promoters also demonstrates the general prevalence of transcription initiation events within gene bodies as observed elsewhere (Carninci *et al*, 2006), although some might in fact reflect recapping of degradation products (Lenhard *et al*, 2012).

Analysis of the relative organization of transcription units, to isolate potential regulatory elements and co-regulated loci, led to the identification of numerous TSS positions in intergenic and other non-coding regions. Most of these TSS could be clustered in hypothetical, previously unreported non-coding transcriptional units that may carry a regulatory role, as recently suggested for the telomeric lncRNAs during parasite invasion (Broadbent *et al*, 2015). many of which correspond to transcription initiation in an antisense orientation to coding genes. More generally, this configuration was widespread across the genome, regardless of the coding potential of the sense/antisense pairs, as previously observed (Patankar *et al*, 2001; Militello, 2005; López-Barragán *et al*, 2011; Siegel *et al*, 2014). Such arrangements that presumably yield overlapping transcripts may carry a regulatory role, as shown by the expression of certain sense-antisense transcript pairs in a mutually exclusive manner. In multiple loci, antisense events originated from bidirectional transcription activity, which we detected for most of the gene pairs in a head-to-head arrangement. For many, we observed coordinate usage of the TSS in both orientations, indicative of the transcriptional units’ co-expression. However, we also identified genomic loci for which divergent transcriptional events were asynchronous or sometimes mutually exclusive, suggesting the possibility of stage-specific regulation of the directionality of bidirectional promoters. Altogether these observations point towards the probable presence of bidirectional promoters with shared and separate regulatory elements (Trinklein *et al*, 2004).

In contrast to the majority of genes for which concomitant usage of the associated TSS was detected, a minority appeared to switch from one TSS to another in a stage-specific manner. This indicates that TSS usage may be temporally controlled, resulting in the generation of developmentally regulated variants. This observation suggests the existence of a context-dependent selection of the TSS, whereby regulatory processes guide the choice for alternative promoters. Interestingly, dynamic TSS usage has been reported in the context of tissue-specificity (FANTOM Consortium and the RIKEN PMI and CLST (DGT) *et al*, 2014) and embryonic development (Haberle *et al*, 2014) and attributed to the differential activity of chromatin-defined enhancers (Andersson *et al*, 2014), or a switch between TSS selection mechanisms (Haberle *et al*, 2014). While several studies argue for the existence of cis-regulatory sequences in the genome of *P. falciparum* (Horrocks *et al*, 2009), further investigations will be needed to assess whether these or other promoter switching mechanisms influence this choice. Local changes in nucleosome configuration mediated by distinct nucleotide compositions (Haberle *et al*, 2014), altered chromatin states (Davuluri *et al*, 2008) or the binding of regulatory factors such as the ApiAP2 transcription factors to specific regions around the TSS (De Silva *et al*, 2008; Campbell *et al*, 2010) may for instance be at play. Given the restricted number of *P. falciparum* genes that display such a behavior of dynamic TSS usage, it will also be interesting to investigate whether this particular mode of transcription regulation is linked to the biological function of these genes. Indeed, the developmentally regulated promoter switch for this subset of genes may be a mechanism to transcriptionally control the stability or translation efficiency of the associated transcripts, or even their biological function, when it is needed the most.

With more than 60% of *P. falciparum* genes for which no biological function has been assigned (Brehelin *et al*, 2008), assessing the functional consequence of a developmentally regulated switch between TSS will require further mechanistic studies.

Our precise mapping of TSS genome-wide revealed an unexpectedly complex and dynamic transcriptomic landscape and constitutes a major advance towards deciphering the molecular basis of *P. falciparum* transcriptional control. Our analysis of the core promoters’ structure notably indicates the existence of a TSS signature despite the exceptionally AT-biased nucleotide composition of the parasite genome. We postulate that spatial constraints mediated by local base composition and nucleosome occupancy may control the accessibility of the basal transcriptional machinery to promoter regions, and thus frequency of TSS usage. This highly valuable resource therefore opens new avenues into the characterization of regulatory elements and the mechanisms directing TSS selection. The recent implementation of CRISPR/Cas9 approaches to edit the parasite genome (Ghorbal *et al*, 2014; Wagner *et al*, 2014) will permit further examination of the interactions between promoters and transcription factors, as well as whether widespread non-coding transcriptional activity plays a role in regulating gene expression.

## Material and Methods

### Parasite culture

The laboratory reference *P. falciparum* strain 3D7 was propagated under standard conditions (Trager & Jensen, 1976). Parasites were synchronized by two consecutive treatments with 5% sorbitol for three or more successive generations before initiating time point samplings every 8 hours throughout the IDC.

### 5’ ends capture and library preparation

Details of sample preparation, 5’ cap sequencing (Pelechano *et al*, 2015) and data analysis can be found in the Supplementary Material.

In brief, single-strand ligation to transcripts’ 5’ end was performed using an RNA adapter containing a 8-mer molecular barcode. cDNA second-strand was synthesized using a biotinylated primer complementary to the RNA 5’ adapter. All samples were treated in the presence of in vitro transcripts (Supplementary Fig S12). Biotinylated samples were captured using streptavidin-coupled Dynabeads M-280 (Invitrogen) and double-stranded cDNA libraries were amplified using KAPA HiFi HotStart ReadyMix (KAPA Biosystems).

Libraries were sequenced on an Illumina HiSeq-2000 (105 bp paired-end sequencing) and reads were mapped to the 3D7 reference genome using GSNAP. Uniquely mapped tags were kept and molecular barcode information was used to filter out PCR duplicates. Processed data and annotations were visualized in an IGB browser (Nicol *et al*, 2009).

### Northern Blot analysis

Probe templates were amplified from 3D7 genomic DNA and labeled using DIG Northern Starter Kit (Roche) as per manufacturer’s instructions. Hybridizations were performed at 55°C. Detailed description can be found in the Supplementary Material.

### TSS annotation, antisense and bidirectional TSS detection

To define TSS blocks morphological operations (Heijmans *et al*, 1989) were performed on the tag counts that were summed across time points and replicates for each position of the genome.

We retrieved the genomic coordinates of every block associated with the optimal filtering step of each genomic feature to generate the final annotation file. The HTSeq package (Anders *et al*, 2014) was used to evaluate the number of 5’ ends falling within an annotated gene feature or a newly annotated TSS block. TSS blocks identified on the opposite strand of annotated features were computed within a window of 500-bp around their 3’ end to detect antisense transcription initiation events. Pairs of annotated features in a divergent arrangement that were separated by a distance of 1-kb or less (Trinklein *et al*, 2004) were analyzed for possible bidirectional transcriptional activity.

### Analysis of the core promoter architecture

To analyze the chromatin organization around the TSS, we used ChIP-seq datasets for H3K9ac, H3K4me3, H2A.Z and H2B.Z from (Bártfai *et al*, 2010) and (Hoeijmakers *et al*, 2013b), respectively. For each gene with at least one annotated TSS block, we used bins of 10-bp within a 2500-bp window (from -1000 to +1500-bp) around the start of the block with the highest counts and estimated the coverage in the genomic locations overlapping these bins. Counts were then summed across all stage-specific genes and divided by the total number of reads mapped in each sample. The occupancy ratio for each histone mark or variant correspond to the ratio between the counts in the IP sample and the chromatin input.

For each gene with at least one annotated TSS block, we computed the local GC content in 10-bp bins around the start of the block with the highest counts. To identify potential nucleotide biases at the site of transcription initiation, we isolated for each gene the TSS block with the highest average counts along the cycle and extracted the nucleotide content in a 5-bp window around the genomic coordinate corresponding to the highest peak of collapsed 5' tags.

### Analysis of the dynamics of TSS usage

A likelihood ratio test was performed to identify TSS blocks for which the usage was different for at least one of the six time points in both biological replicates. TSS blocks with a local FDR < 0.01 were considered active. Transcript level at every single time point was tested against the mean value calculated across all time points using DESeq2 (Love *et al*, 2014). An absolute log2 fold change ≥ 0.5 was required to identify peaking time points.

Details on the identification of transcript pairs with highly distinct TSS usage patterns are described in the Supplementary Material.

## Data access

All 5’ cap sequencing raw data from this study are accessible through the GEO accession number GSE68982.

## Acknowledgements

We thank Emilie Fritsch and Marcus Lee for helpful comments on the manuscript, and Aleksandra Pekowska for fruitful discussions over the course of the project.

We thank Richard Eastman for kindly providing the *P. falciparum* strain 3D7. This study was technologically supported by the EMBL Genomics Core Facilities. S.H.A was supported by an EIPOD / Marie Curie COFUND postdoctoral fellowship. C.D.C was supported by a PhD fellowship from the Boehringer Ingelheim Fonds. This study was supported by the National Institutes of Health Grant NIH R01 GM068717 (to L.M.S.).

## Author contributions

S.H.A., C.D.C, V.P. conceived the study. S.H.A. performed the experiments with contributions from C.D.C. and V.P. S.H.A., C.D.C., and B.K. performed the analysis. C.D.C. developed the morphological mathematics approach for the *P. falciparum* TSS data. C.D.C. and B. K. established the statistical pipeline for the dynamic analysis. S.H.A. and C.D.C. interpreted the results, together with V.P. and L.M.S. S.H.A and C.D.C wrote the manuscript, with inputs from the other contributors. All authors read and approved the final manuscript.

